# A Spatial Multi-Modal Dissection of Host-Microbiome Interactions within the Colitis Tissue Microenvironment

**DOI:** 10.1101/2024.03.04.583400

**Authors:** Bokai Zhu, Yunhao Bai, Yao Yu Yeo, Xiaowei Lu, Xavier Rovira-Clavé, Han Chen, Jason Yeung, Georg K. Gerber, Mike Angelo, Alex K. Shalek, Garry P. Nolan, Sizun Jiang

## Abstract

The intricate and dynamic interactions between the host immune system and its microbiome constituents undergo dynamic shifts in response to perturbations to the intestinal tissue environment. Our ability to study these events on the systems level is significantly limited by *in situ* approaches capable of generating simultaneous insights from both host and microbial communities. Here, we introduce <u>Micro</u>biome <u>Cart</u>ography (MicroCart), a framework for simultaneous *in situ* probing of host features and its microbiome across multiple spatial modalities. We demonstrate MicroCart by comprehensively investigating the alterations in both gut host and microbiome components in a murine model of colitis by coupling MicroCart with spatial proteomics, transcriptomics, and glycomics platforms. Our findings reveal a global but systematic transformation in tissue immune responses, encompassing tissue-level remodeling in response to host immune and epithelial cell state perturbations, and bacterial population shifts, localized inflammatory responses, and metabolic process alterations during colitis. MicroCart enables a deep investigation of the intricate interplay between the host tissue and its microbiome with spatial multiomics.

## Introduction

The intestinal environment represents a highly intricate ecosystem characterized by diverse and dynamic interactions, including the mucosal layer and its plethora of bacterial components (1), and the immune and epithelial cell populations within its adjacent tissue (2). The balanced interplay between these microbiome components, epithelial cells, and immune players is crucial for maintaining immune homeostasis (3). The remarkable ability of microbial species to modulate and educate the host immune system starts early during infancy (4). This is achieved via the delivery of antigens to gut resident T cells and other immune populations, which in turn promotes immune tolerance and plays a vital role in immune homeostasis in humans (5). Recent studies have also unveiled the immune-modulatory effects of the microbiome in tumor patients, wherein specific microbiome-related peptides can cross-activate tumor-infiltrating lymphocytes, thereby facilitating more effective patient treatments (6, 7). When this delicate balance of the host-microbiome homeostasis is disrupted, particularly in the presence of various external perturbations or disease states, a myriad of other interactions emerge in response to physical barrier breaches. For instance, in inflammatory bowel disease (IBD), functional changes are observed in epithelial, immune, and bacterial cells, leading to a compromised segregation among these elements, and eliciting acute immune responses in the intestinal tissue (8). Colorectal cancer represents another prominent example wherein altered interactions within the intestine during disease states contribute to the deterioration of the epithelial cell layer and microbiome dysbiosis (9). This, in turn, results in increased delivery of bacterial toxins to the tissue, causing DNA damage and exacerbating tumor progression. Overall, understanding the intricate interplay between the microbiome, epithelium, and immune cells within the intestinal tissue microenvironment is of utmost importance in comprehending the maintenance of a healthy system as well as the pathological mechanisms underlying various diseases.

Current approaches for investigating these interactions, whether in a homeostatic or diseased state, present significant challenges. Traditional tools, such as 16S rRNA sequencing or metagenomic sequencing for analyzing the microbiome, and flow cytometry or single-cell RNA sequencing (scRNA-seq) for studying host cells, have provided valuable insights into different populations in various settings. Although these methods offer in-depth information, they often lack crucial spatial context, missing the unique opportunity to study interaction events in their native environment. Fortunately, a growing number of methods have emerged in recent years, aiming to decode host cell populations and functions with spatial information. These approaches include multiplexed imaging techniques (10–12) and spatially resolved sequencing methods (13–15). These innovative techniques enable the examination of host cell interactions within their specific locations. Similarly, advances have been made in understanding the spatial organization of the microbiome within its native context. Initially, methods were developed to target specific bacterial groups (16), but more recent approaches have enabled the investigation of hundreds of bacterial species (17, 18) using cyclic imaging or modified sequencing assays. Multiplexing 16S-specific probes with human poly-A capture in spatially resolved sequencing “spots” has also been a significant advancement in spatially resolving host-bacterial interactions in situ (19, 20). These approaches have greatly facilitated our understanding of the intricate interaction between host cells and microbial components from a spatial perspective. Additional spatial multiomics approaches are required for a more comprehensive understanding of host-microbial interactions, including 1) bacterial identity, to study microbiome spatial patterns; 2) protein expression, particularly for defining immune-epithelial communities and related niches; 3) transcriptomic, to elucidate the functional shift in presence of spatial change in microbiome, immune, and epithelial communities; 4) glycomics, to investigate the metabolic change in response to functional alterations.

To address this technological gap, we introduce <u>Micro</u>biome <u>Car</u>tograph (MicroCart), an integrative framework designed to bridge the divide between host-microbiome interactions and spatial analysis. MicroCart consists of an optimized 16S probe design and validation approach for highly specific targeting of bacterial taxa, while also ensuring the co-preservation of diverse biological targets within the tissue for downstream investigations using multiplex imaging platforms (Multiplexed Ion Beam Imaging; MIBI), spatial sequencing modalities (Nanostring GeoMx Digital Spatial Profiling; DSP), and mass spectrometry imaging techniques (MALDI-IMS imaging for N-glycans), thus enabling comprehensive and multi-omics spatial dissection of host-microbiome interactions across any biological model.

## Results

### Overall study design for MicroCart

The MicroCart framework allows a detailed investigation into tissues of interest that contain intact microbial components, such as intestine tissues from a mouse model of colitis as performed in this study (**Fig. 1A**): In brief, we induced colitis in mice by introducing 3.5% DSS in drinking water for 6 days. A total of 4 mice were included in the colitis group, while 4 mice served as a healthy control group with normal drinking water. The intestinal tissues collected from the mouse experiment can be subject to various spatial-omics techniques, including the simultaneous probing of 1) MicroCart-MIBI imaging of both microbial and host components with antibodies and 16S-specific probes (**Fig. 1A, top**), 2) MicroCART-GeoMx spatial transcriptomics with custom 16S-specific probes, in conjunction with murine whole transcriptome-level capabilities (13) (**Fig. 1A, middle**), and 3) MALDI-MSI for N-glycans using mass spectrometry (**Fig. 1A, bottom**) on serial sections from the same tissue for a tri-modality spatial interrogation of host-microbe interactions *in situ* (**Fig. 1B**).

**Figure 1:**
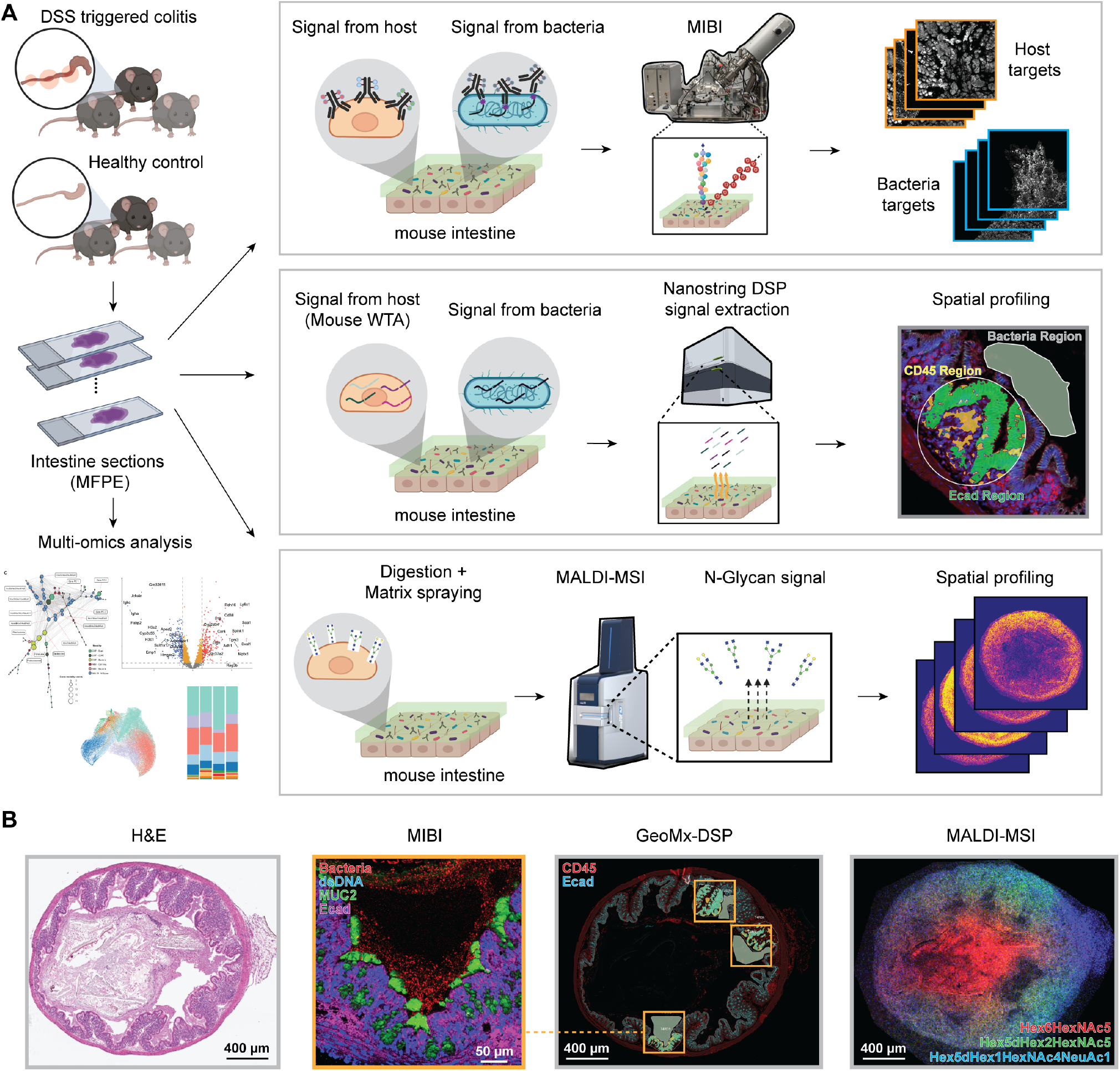
(A) Overview of the study. Colitis mice (triggered by DSS, n = 4) and healthy control mice (n = 4) were sacrificed, and intestinal tissues were dissected. Tissue samples were fixed with the Methacarn-formalin method developed here and embedded in paraffin (MFPE; detailed in Material & Methods), before sectioning onto serial slides. These adjacent slides were subject to analysis using: 1) Multiplex Ion Beam Imaging (MIBI) spatial proteomics, with an antibody panel targeting host antigens and custom oligo probes targeting bacterial groups; 2) GeoMx Digital Spatial Profiler (DSP) spatial transcriptomics, with a whole transcriptome panel targeting host RNA molecules and custom oligo probes targeting bacterial groups. The selected DSP regions are aligned to where the MIBI FOVs were acquired within the adjacent slides; 3) MALDI mass spectrometry imaging (MALDI-MSI) that measures N-Glycan levels. **(B)** Representative images of the intestinal tissue sections that were investigated by the three different modalities. Images from left to right: H&E image of the tissue section; representative MIBI antibody signals from the tissue section; fluorescence image with boxes indicating the regions being captured for transcriptomic analysis in the tissue section of DSP; representative MALDI N-Glycan signals from the tissue section.

### Robust design and efficient validation of bacteria oligo probes

The imaging of bacteria using *in situ* hybridization (ISH)-based methods has been a longstanding approach in the field (21). To achieve this, oligonucleotide probes specifically targeting conserved 16S ribosomal RNA sequences are first designed *in silico*, enabling the visualization of various bacterial groups (22). Given that most 16S ISH probes currently in use were designed before the advent of next-generation sequencing (NGS) technologies (23), a significant number of existing ISH probes in the literature may not accurately target the intended bacterial groups (24). To address this limitation, we first introduce an improved 16S ISH probe designing pipeline that aims to achieve robust and precise targeting of bacteria, within the context of the intact intestinal microbiome (**Fig. 2A**). A major bottleneck for probe design is the delicate balance between the coverage and specificity of the probes, as demonstrated for eukaryotic genomes (25). This process is even more challenging in the microbial context, considering the vast amount of bacteria sequences publically available in conventional databases (26, 27). To overcome this hurdle, we adopted a strategy (28) that involved constructing a curated 16S rRNA sequence pool exclusively consisting of known bacteria found in the intestinal microbiome, totaling 12,936 near-full length 16S rRNA sequences. We then employed phylogeny sequence searcher ARB (29) to identify signature sequences from this curated sequence pool that qualify for the coverage and specificity requirements for a user-defined bacteria target group. Subsequently, these candidate signature sequences as identified by ARB are subjected to additional filtering and screening based on multiple criterion, including melting temperature, hybridization efficiency (30), and predicted secondary structure (31). Probes that meet all the above criteria are retained. To target groups at lower phylogeny levels, such as the species level, a single oligonucleotide probe is used. However, for higher phylogeny levels, like the phylum level, a single probe often fails to provide satisfactory coverage and specificity due to the large number of sequences within the target group. Therefore, we employed an additional combinatory probe set strategy, where multiple probes selected from the previous step were combined into groups of three. These probe combinations were then evaluated *in-silico* for optimal coverage and specificity, where fast sequence alignment of probes to the curated 16S rRNA sequence pool with Usearch (32) was performed. Following this step, the probe combination group that achieved the highest coverage while maintaining specificity is then recommended for experimental validation (**Fig. 2A**). This probe design component of MicroCART is capable of rapidly designing highly specific 16S probes against microbiome components at various levels (**Supp. Table 1**).

**Figure 2:**
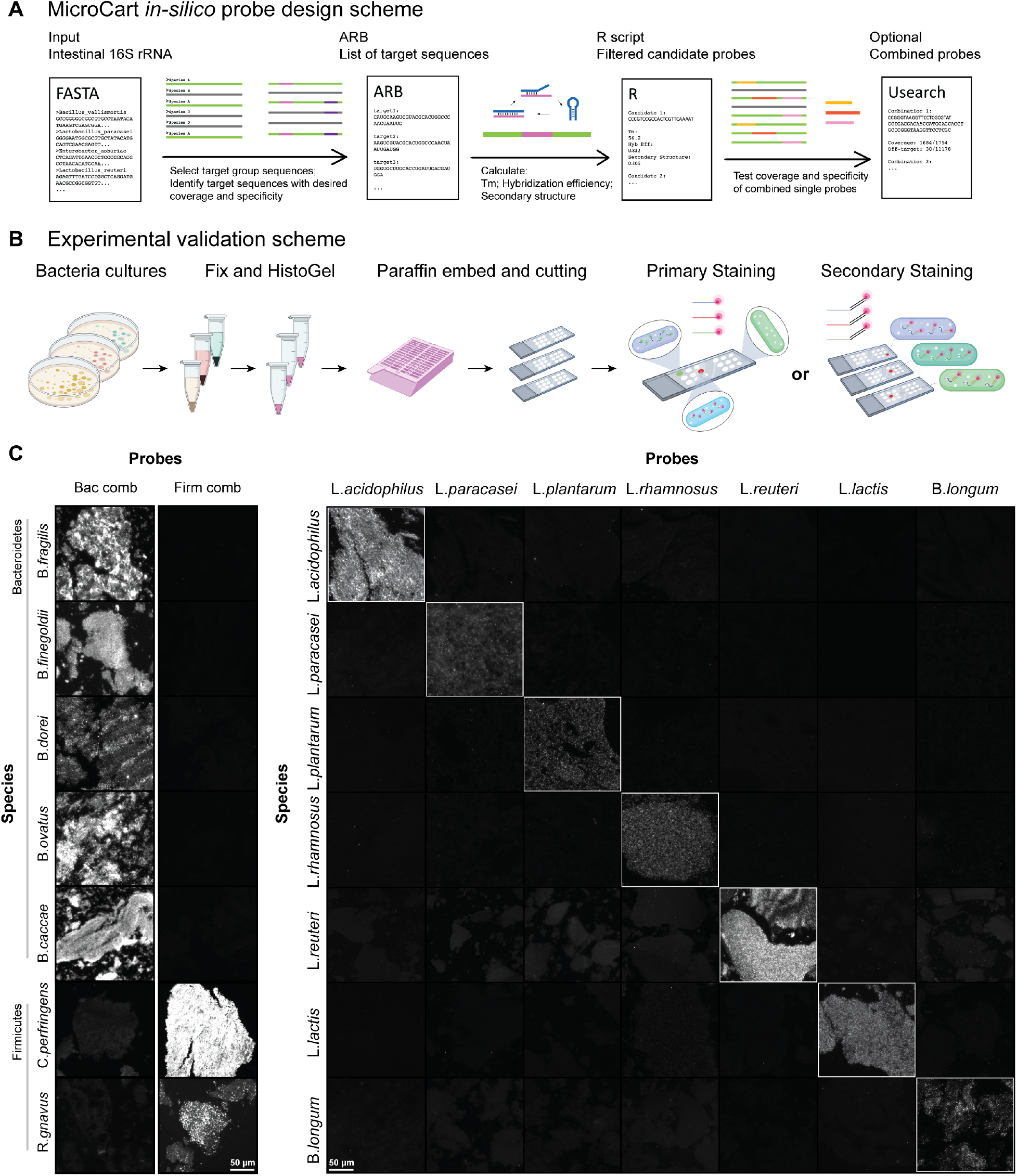
(A) Schematic of the MicroCart *in silico* probe design process. A curated database of 16S rRNA sequences from intestinal microbiota was first created, then probe candidates were designed that are specific for target bacterial groups. Stringent criteria, including melting temperature, hybridization efficiency, secondary structure, coverage, and specificity were used to select for top candidate probes. Optionally, the MicroCart probe design tool can also create combination sets of probes to maximize performance. **(B)** An illustration of the experimental validation process for designed probes. Bacteria strains were cultured, harvested, fixed (using our optimized MFPE fixation), and embedded in HistoGel. Subsequently, HistoGel-bacteria strains were dehydrated and embedded in paraffin in a microarray fashion, into Bacteria MicroArrays (BMA), before sectioning onto slides. Probe validation can be either performed using primary oligos conjugated to fluorophores, or using a secondary barcoded-oligos conjugated to fluorescence to reduce cost through flexibility and increase efficiency through multiplexing. **(C)** Left: Experimental validation on BMA slides, with probes designed to target phylum groups (Bacteroidetes or Firmicutes). Right: probes designed to target various probiotic species.

Coupling the *in silico* pipeline of MicroCart with a rigorous experimental validation framework is crucial for ensuring the quality, specificity, and reliability of the probes. Traditional methods for validating bacteria ISH probes involve growing bacterial cultures, and performing brightfield or fluorescence ISH staining using labeled candidate probes. However, this process is not easily scalable, and does not include proper controls to assess probe specificity. To address this challenge, we introduce an efficient bacteria probe validation pipeline in MicroCart (**Fig. 2B**). We first culture multiple related and non-related intestinal microbiome bacteria species (**Fig. 2B, left**), with each species being grown in its specific required medium and under anaerobic conditions as needed. After cultivation, the bacteria were harvested and centrifuged to obtain bacterial pellets. The bacterial pellets were then fixed using methacarn and subsequently fixed in methacarn and formalin. To further maintain the structural integrity of the fixed bacteria pellets, we embedded them in Histogel (33). The fixed and histogel-embedded bacteria pellets across different species were finally arranged in an array format in a tissue cassette, and embedded in paraffin to create a methacarn and formalin-fixed, paraffin embedded (MFPE) Bacteria MicroArray (BMA). The MFPE-BMA allows for repeated sectioning and analysis, eliminating the need for repetitive bacteria culturing whilst maintaining the fixation conditions as the final tissues of interest for maximal compatibility. By using the MFPE-BMA slides, probes can either be validated with the conventional primary probe with fluorescence, or a primary and secondary oligo staining scheme akin to Oligopaints for cost efficiency (34), where all the candidate probes that undergo testing can share a conserved secondary detection barcode, thus reducing the required amount of fluorescence-labeled detection probes (**Fig. 2B, right**). Most importantly, these control MFPE-BMA sections contains multiple bacteria species simultaneously, allowing for a combined positive and negative control for ISH specificity and assay performance. Using this improved design and validation pipeline for bacteria probes in MicroCart, we were able to design and validate probes targeting different phylogeny levels of groups. At the phylum-level, we designed probes targeting the phylum Firmicutes and Bacteroidetes (**Fig. 2C, left**). In parallel, we also designed and validated single oligonucleotide probes targeting various species-level probiotic taxa, including: Lactobacillus acidophilus, Lactobacillus paracasei, Lactobacillus reuteri, Lactococcus lactis, Lactobacillus plantarum, Lactobacillus rhamnosus, and Bifidobacterium longum subsp. longum (**Fig. 2C, right**). Probes designed using the MicroCart pipeline performed robustly on both levels, with fluorescent signals observed only in targeted groups but not the others, and limited off-targeting binding in bacteria with close sequence similarity (**Supp. Fig. 1, Supp. Table 2**).

### Multiplexed imaging of microbiome and host cells with MicroCart-MIBI

We next adapted the probes produced with our MicroCart pipeline onto multiplexed imaging platforms, as exemplified with the MIBI-TOF, an imaging platform capable of >40-plex spatial readout using secondary ion mass spectrometry combined with a time-of-flight readout to resolve metal-tagged labels in tissue sections at subcellular resolutions (35). To further amplify the oligo targets for an optimal signal-to-noise ratio, we adapted the metal conjugatedantibody-based approach by targeting labeled antibodies specific for haptens covalently tagged to 16S-targeting oligos, to achieve signal amplification beyond standard metal-tagged oligos (36, 37). This allowed for robust detection of bacterial signal on the MIBI-TOF in conjunction with antibodies against haptens of interest (**Supp. Fig. 2**). To accurately localize the spatial patterns of both microbiome and host components, we also further improved upon the fixation method for microbiome-related tissues. Methacarn fixation is commonly suggested for the preservation of mucus structure and bacteria localization in tissue samples (38), but this approach is not as effective for the preservation of protein epitopes as formalin fixation and paraffin embedding (FFPE), the current clinical standard for tissue preservation widely used in standard clinical histology, and spatial platforms including the MIBI-TOF (39–41). We postulated that an optimized fixation method suitable for MicroCart should ideally contain both fixatives: first methacarn to preserve the mucosal layer and the bacterial components within (38), followed by formalin to ideally preserve protein epitopes. This approach, termed Methacarn and Formalin-fixed, Paraffin-Embedded (MFPE), was validated for it’s ability to preserve both the mucosal layer and protein epitopes (**Supp. Fig. 3& 4**).

**Figure 3:**
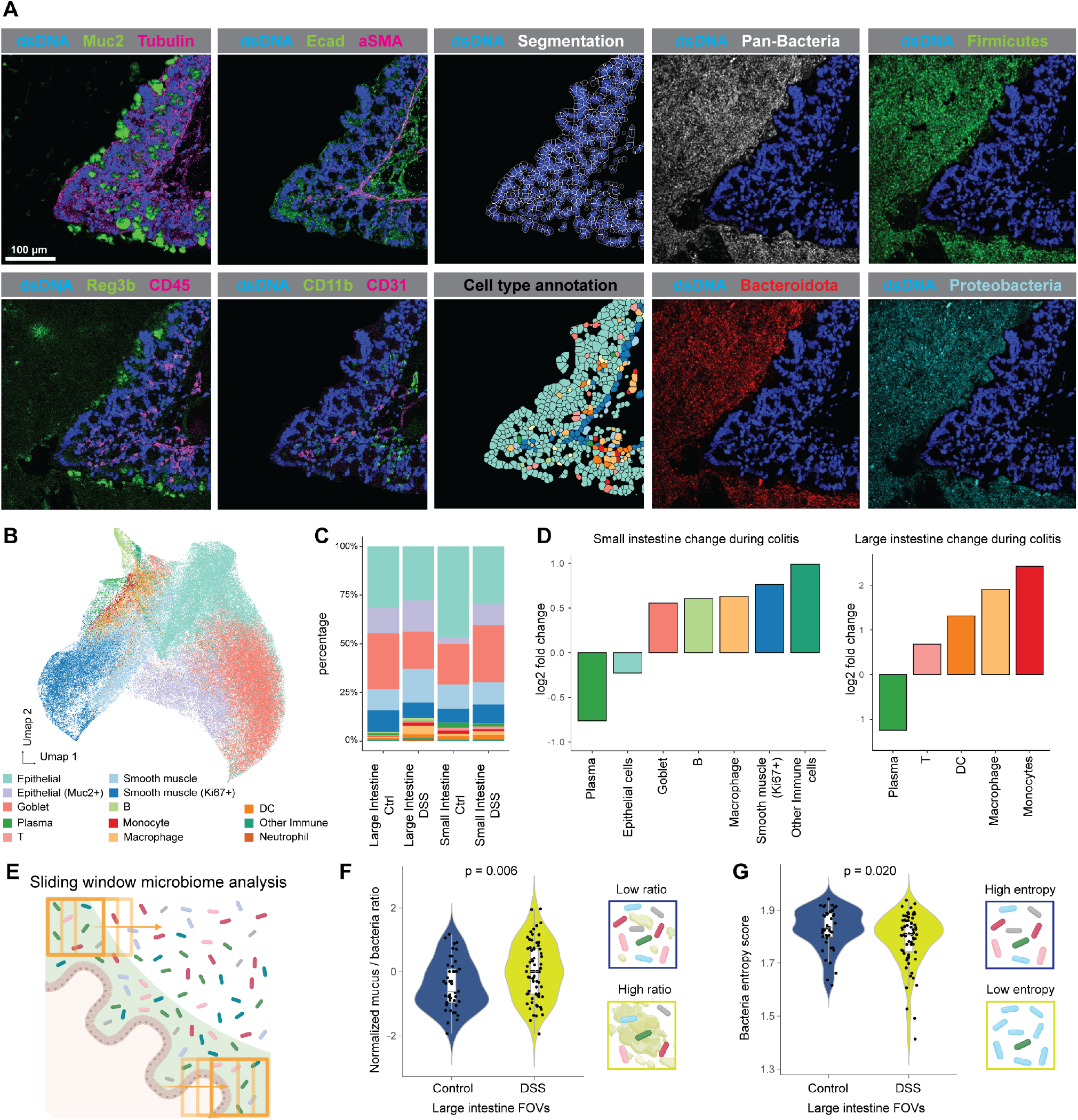
(A) Images from a representative tissue region showing selected MIBI signals, cell segmentation, and cell type annotation information from host cells or microbiome communities. **(B)** UMAP dimensional reduction visualization of host cell type annotation information, based on single cell MIBI protein expression profiles. **(C)** Cell type proportions per tissue sample, grouped by tissue location and colitis status. **(D)** Cell types with a significant (p.adj < 0.05, Student’s t-test) frequency change compared between colitis and healthy tissues. Left: small intestine. Right: large intestine. **(E)** Illustration of the sliding window microbiome analysis devised for the MIBI microbiome analysis in (F) and (G). **(F)** Violin plot and illustration of the localized mucus-bacteria ratios in control or colitis large intestine tissue samples, p-value calculated with Student’s t-test. For more details, see Material & Methods. **(G)** Violin plot and illustration of the localized bacteria Shannon entropy in control or colitis large intestine tissue samples, p-value calculated with Student’s t-test. For more details, see Material & Methods.

MFPE tissue sections were first subject to ISH with MicroCart-oligos carrying covalently attached haptens (**Supp. Table 3**), followed by staining with an MIBI antibody cocktail panel including antibodies that bind specifically for these haptens (**Supp. Table 4**), thus enabling simultaneous imaging of both host proteins and bacterial components (**Fig. 3A, Supp. Fig. 5A**). We performed MIBI imaging of a total of 202 field-of-views (400um * 400um; FOVs), across intestinal tissues from DSS-treated and healthy control mice (n = 4 each), identifying a total of 126,426 host cells, inclusive of diverse immune and epithelial cell types within the small and large intestines (**Fig. 3B, Supp. Fig. 5B**). We observed significant compositional changes in the intestinal cell populations during colitis (**Figs. 3C & D**), including reduced numbers of plasma cells in both small and large intestines in DSS-induced colitis, and a global increase in immune cells in both the small and large intestines, reflective of the localized nature of the immune response to colitis.

**Figure 4:**
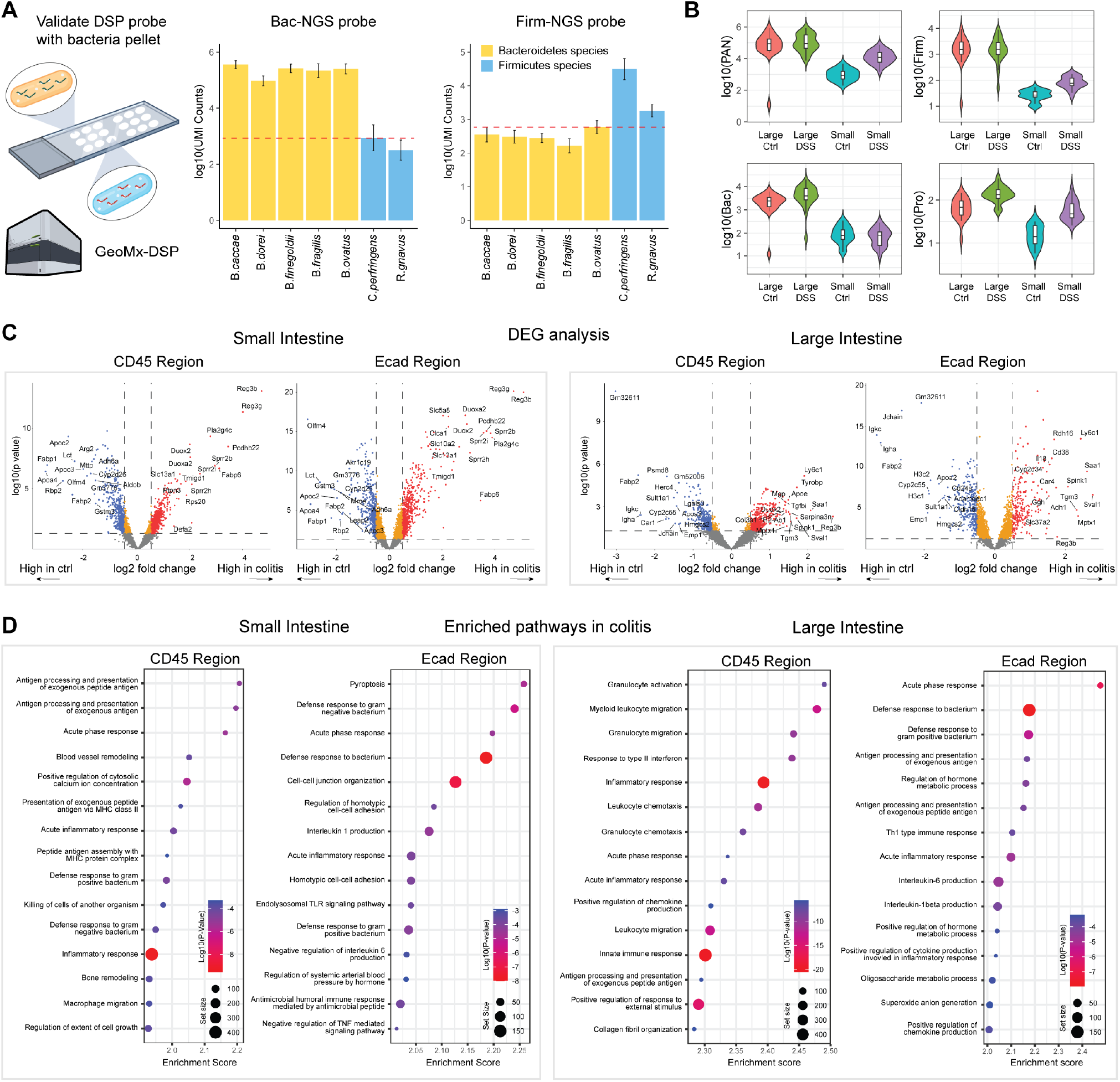
(A) Left: Illustration of the validation scheme for bacteria probes on the BMAs, using MicroCart-DSP spatial sequencing as a read out. Right: Barplots showing sequencing counts (mean ± 0.95 CI) from the respective probes in different bacteria species arrays. Red dotted line indicates the limit of detection. **(B)** Violin plot showing bacteria probe counts from MicroCart-DSP coupled with the mouse Whole Transcriptome Atlas probes, summed up from individual bacterial regions in tissue sections, separated by colitis status and tissue locations. **(C)** Volcano plots showing top differentially expressed genes for host cells between healthy and colitis samples, separated by MicroCart-DSP region compartments (E-Cad+ or CD45+) and tissue locations. **(D)** GSEA analysis showing the top 15 enriched gene pathways in the colitis groups, separated by MicroCart-DSP region types and tissue locations.

**Figure 5:**
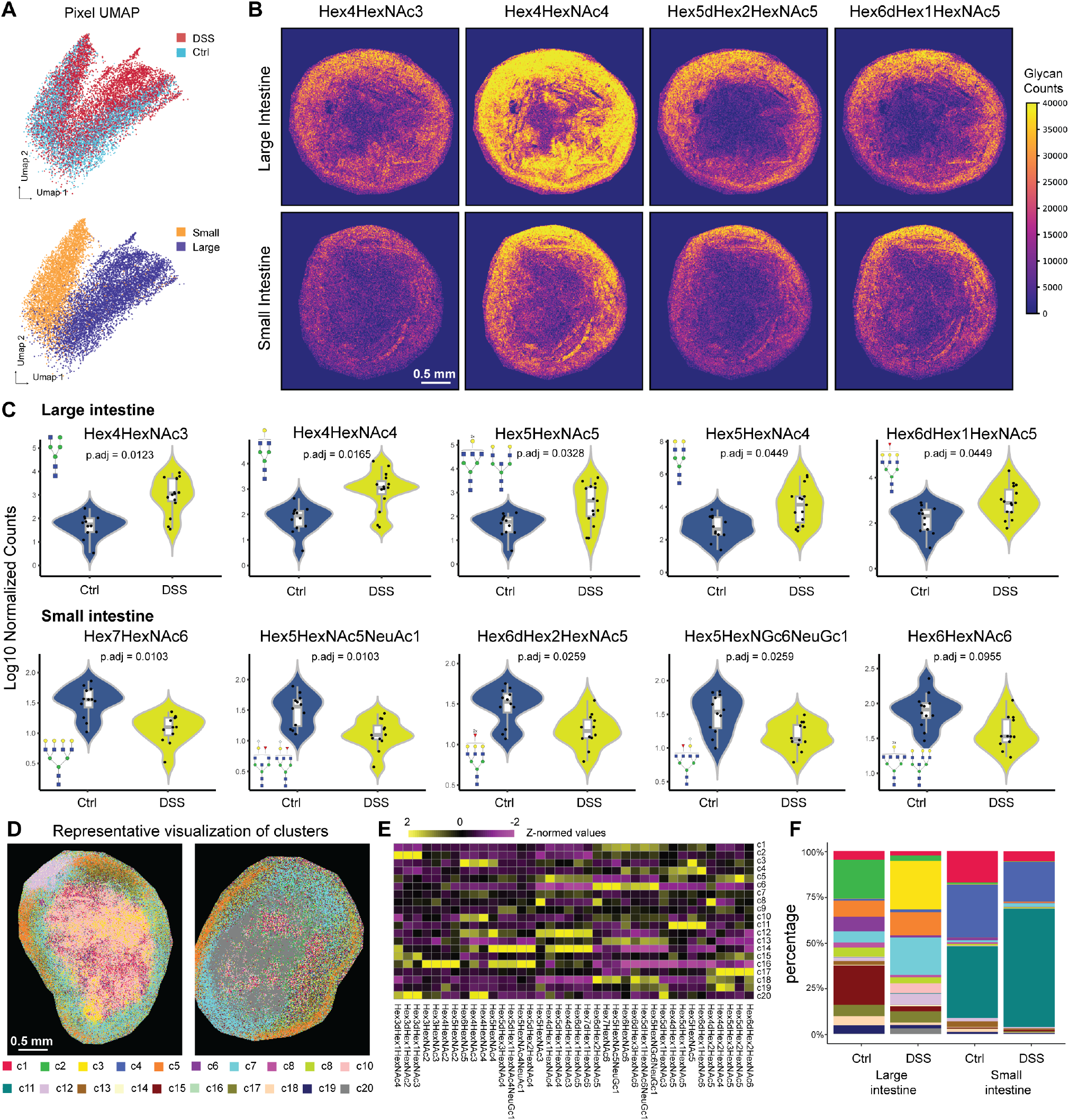
(A) UMAP dimensionality reduction visualization of colitis status or tissue type information, based on pixel level N-glycan signals from MALDI-MSI. Pixels were either colored by colitis status or tissue location. **(B)** Representative images of selected N-glycan signals from small and large intestinal tissues. A total number of 51 intestinal tissue sections (all tissue sections that were investigated by MIBI and DSP) were imaged using MALDI-MSI. **(C)** Top 5 most significantly changed (p.adj < 0.05, Student’s t test) N-glycans in small and large intestine tissues compared between healthy and colitis status. **(D)** Representative images of pixie (49) clustering results based on N-glycan signals. **(E)** Heatmap of the average N-glycan level for each pixel level cluster from pixie. **(F)** Pixie cluster percentage for each tissue across colitis status and tissue locations.

We next quantified the spatial variations in the bacterial components during colitis using a sliding window strategy, focusing on quantitative spatial variations across the intestinal FOVs (**Fig. 3E**). We first evaluated the ratio of mucosal size to bacteria patch within each sliding window, among the non-host region in each MIBI FOV. A higher value indicated increased local intermixing between the host mucus and bacterial community (see Material & Methods for more details). Our results indicated that mice with colitis exhibited significantly more local intermixing of host and bacteria cells, reflective of potential microbial-linked remodeling and barrier penetration related to DSS-induced colitis (**Fig. 3F**). We next assessed the local Shannon entropy of various bacterial phyla based on our MicroCart ISH probes, wherein a lower entropy value reflected decreased diversity in the local microbiome composition, as was observed in mice with colitis (**Fig. 3G**). Together, these results highlight the ability of MicroCart, coupled with highly multiplexed imaging, for a multi-modal dissection of the host-microbial remodeling and spatial reorganization in a mouse-model of colitis.

### Spatially resolved sequencing of host and microbiome with MicroCart-DSP

To orthogonally validate our bacteria probe specificity (**Fig. 2C**), we developed a customized workflow (see Material & Methods) for the spatial transcriptomics Nanostring GeoMx DSP platform, using MicroCart-DSP custom probes (**Supp. Table 5**) on MFPE-BMA sections. We successfully confirmed the specificity of MicroCart-DSP probes to their targeted phylum group using NGS sequencing as a readout for the unique UMIs on these MicroCart barcodes (**Fig. 4A**), highlighting the cross-platform utility of MicroCart designed probes for both spatial imaging and sequencing. Given the feasibility of MicroCart-DSP custom probes *in situ*, we next developed a custom workflow for integrating the mouse whole transcriptome atlas (WTA; > 20,000 genes) probeset in conjunction with MicroCart-DSP custom probes, to spatially dissect cellular pathways, immune signaling, metabolic states and microbial compositional changes between healthy and colitis mouse intestines, as stratified by CD45-positive (immune), E-cadherin-positive (non-immune epithelial), and bacterial regions. We used this approach to sequence a total of 350 regions. Akin to our MFPE-BMA results, our mouse tissue spatial transcriptomes results also confirmed the specificity of bacterial signals (**Fig. 4B, Supp. Fig. 6**). This MicroCart-DSP approach enabled the further investigation of the host-pathogen responses during colitis. In the small intestine regions, we observed an increase in the expression of genes in the Reg3 family (*Reg3b & Reg3g*), both previously implicated with potent roles in antimicrobial activity and tissue repair during colitis (42). We also observed an increase in gene expression of isoenzymes *Duox2* and *Duoxa2*, previously implicated with IBD in human patients (43, 44), and the Sprr2 family (*Sprr2b, Sprr2i & Sprr2h*), suggestive of an antimicrobial response specific for Grampositive bacteria (45, 46), signaling a disruption of the bacterial community composition within the small intestines during colitis. Meanwhile, in the large intestine, we observed increased expression of genes including the myeloid cell marker *Ly6c1* and decreased expression of plasma cellrelated genes (*Igkc, Igha & Jchain*), consistent with our MIBI observations on increased macrophages and monocytic infiltrations, and depletion of plasma cells in the large intestine during colitis (**Fig. 3D**). The increased expression of gene *Saa1* further highlights the critical role of macrophages in acute inflammation within the large intestine, in line with studies linking its protein product, Serum Amyloid A, and macrophage infiltration in humans and mice (47, 48).

**Figure 6:**
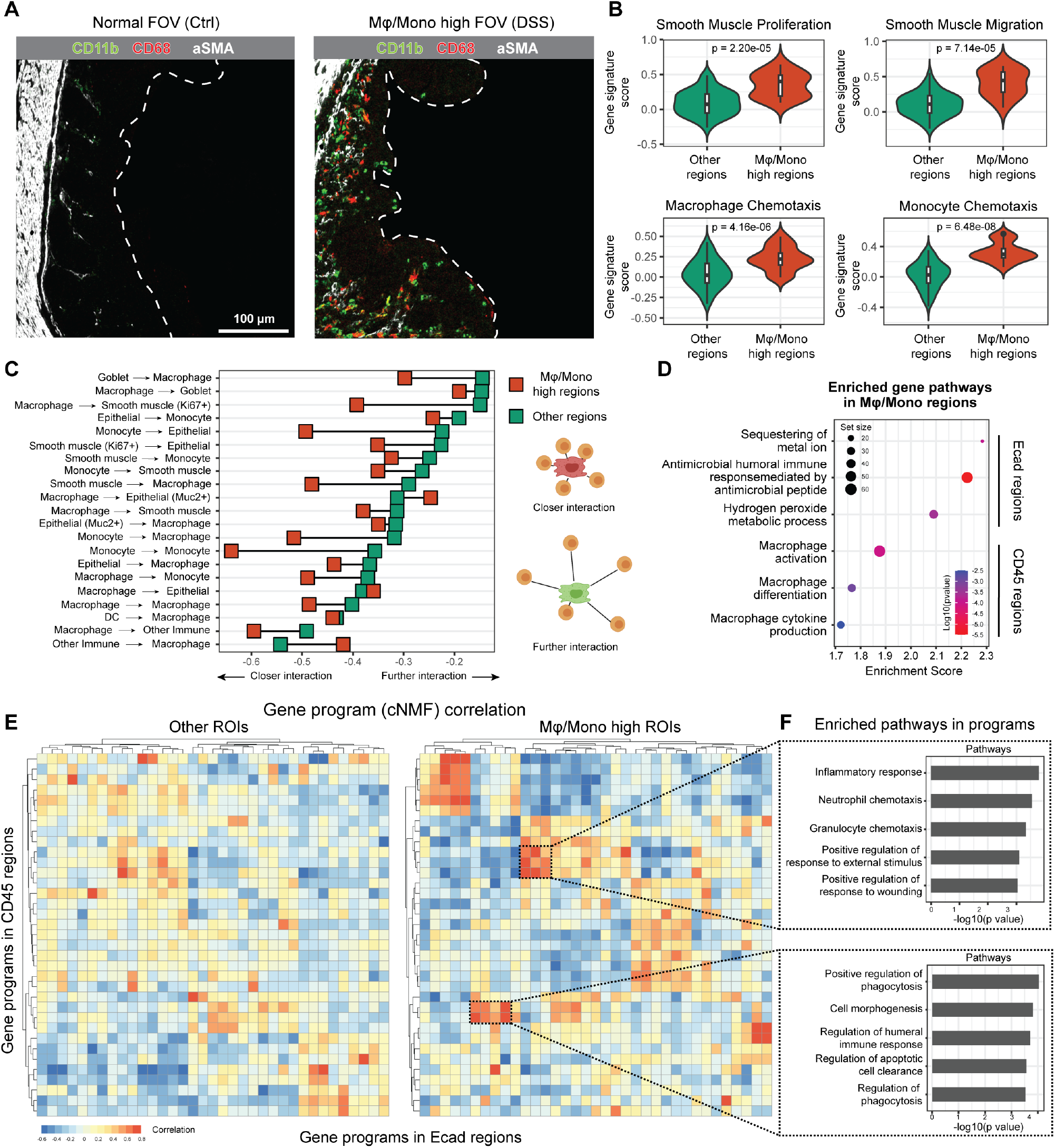
**(A)** Representative MIBI images of a tissue region not infiltrated by macrophages and monocytes (left), and a region with high number of macrophage and monocyte infiltration (right). Only Large intestine regions were considered in this analysis. **(B)** Violin plots of gene pathway scores from the MicroCart-DSP regions, separated by macrophage and monocyte infiltration status (based on the MIBI information from the same regions on the adjacent slide). **(C)** Dumbbell plots of pairwise cell-cell interactions enriched or depleted compared to a randomized background permutations (1000 iterations) background between macrophage/monocytes to other cell types in large intestine tissues, separated by macrophage/monocyte infiltration status. Only interactions that passed a statistical test (p < 0.05, Wilcoxon test) for both infection conditions are shown. **(D)** GSEA analysis on high macrophage/monocyte infiltration regions. For Ecad regions, top 3 enriched significant pathways (p.adj < 0.05) related to anti-bacterial functions were shown. For CD45 regions, top 3 enriched significant pathways (p.adj < 0.05) related to macrophage activities were shown. **(E)** Heatmap of the correlation of gene programs as identified via cNMF (cite) between paired CD45 and Ecad MicroCart-DSP compartments. Left: correlation heatmap from tissues that were not infiltrated by macrophage/monocyte. Right: correlation heatmap from tissues that were infiltrated by macrophage/monocyte. **(F)** Gene ontology (GO) analysis on selected correlation hotspots of gene programs. Top 10 genes that contribute to each of the gene programs within the selected hotspot were grouped together, and used as input for the GO analysis. The top 5 most enriched GO terms were shown for each hotspot.

We next conducted region-specific transcriptomic analysis to better contextualize pathway-level changes in the immune (CD45+) and non-immune (Ecad+) regions during colitis across the mouse small and large intestines. We first performed GSEA on the GeoMx spatial transcriptomic data, and observed the enrichment of pathways related to antigen presentation of exogenous antigens, and the killing of cells from another organism in the CD45+ immune compartment, reflective of an orchestrated immune response to the potential mucosal breach and exposure to bacterial components during colitis. In the Ecad+ region of the small intestine, the enriched pathways in epithelial cells were also predominantly associated with various defensive pathways, including pyroptosis, and others during colitis (**Fig. 4D, left**). Conversely, within the large intestine, the CD45+ immune regions exhibited enrichment in immune responses encompassing pathways related to granulocyte, leukocyte migration, chemotaxis, and activation. In the Ecad+ regions, enriched pathways included immune responses and metabolic processes (**Fig. 4D, right**). These results highlight the diverse range of immune responses and varying metabolic shifts across local regions of the small and large intestines upon DSS-treatment in a mouse model of colitis.

### MALDI-MSI detects global glycosylation shift in intestine during colitis

Given the implications of immune responses and tissue remodeling in response to DSS-induced colitis and microbial changes, we further observed alteration of genes related to the host glycosylation processes (**Supp. Fig. 7A& B**), a key component of immune cell trafficking (50). For example, we found transcripts encoding for glycosyltransferases (*Mgat3, Mgat4a, Mgat4b, Mgat5*) significantly upregulated in the small intestine epithelium layer. We also identified key glycosyltransferases fucosyltransferase (*Fut2*), and galactosyltransferase (*B4galt1*), to be significantly upregulated in both large and small intestine epithelium layers. Lastly, we also observed beta-mannosidase (*Manba*) to be significantly downregulated in both large and small intestine epithelium layers. These genes have also been described as potential IBD risk factors in human patients (51). Given our spatial transcriptomics results, we postulated that tissue glycosylation patterns are linked to the microbial invasion and immune-epithelial remodeling observed in colitis. To comprehensively assess the unknown spatial glycosylation landscape in our mouse colitis model, we implemented timsTOF fleX MALDI-2 N-glycan imaging on the tissue sections adjacent to the ones previously investigated using MicroCart-MIBI and -DSP, to stratify varying N-Glycan tissue components down to a 10 µm pixel-level spatial resolution (**Supp. Table 6**). Dimensional reduction using UMAP on the glycan compositions per pixel stratified between 1) small and large intestines,and 2) healthy and DSS tissues, highlighted both common and unique glycosylation patterns that are species-, tissue- and disease-specific (**Fig. 5A**). We next confirmed these differences from visual inspection of the data across several glycans (**Fig. 5B**). Further quantification of the data identified significant glycosylation changes between healthy and colitis tissues: in large intestine tissues, there was a marked increase in Hex4HexNAc3, Hex4HexNAc4, Hex5HexNAc5,Hex5HexNAc4, and Hex6dHex1HexNAc5 during colitis. Conversely, in small intestine tissues, we observed a notable decrease in Hex7HexNAc6, Hex5HexNAc5NeuGc1,Hex6dHex2HexNAc5, Hex5HexNAc6NeuGc1, and Hex6HexNAc6 during colitis (**Fig. 5C**). Previous studies have found association between intestinal inflammation and upregulated expression of truncated and immature surface glycans (51), which is consistent with the observation in our study, where we detected an increase in low-branching N-glycans and a decrease of high-branching N-glycans in DSS intestinal tissues (**Fig. 5C**). We next employed a pixel-level clustering approach (49) on N-Glycan signals to identify 20 spatially distinctive glycosylation populations (**Fig. 5D &E**). Our results indicate the enrichment of Hex3dHex1HexNAc4, Hex3dHex1HexNAc2, and Hex3dHex1HexNAc3, in the large intestine, as indicated by the significant decrease in clusters 2 (p.adj = 0.00012, Student’s t test), and 15 (p.adj = 0.00235), and increase in cluster 7 (p.adj = 0.00054) during colitis. We further observed in the small intestines during colitis, the enrichment of Hex5HexNAc6NeuGc1, Hex5dHex1HexNAc6NeuGc1, and Hex6dHex3HexNAc6, as represented by the significant decrease in clusters 1 (p.adj = 0.0205) and 6 (p.adj = 0.0205), and increase in cluster 11 (p.adj = 0.0217) (**Fig. 5F**).

**Figure 7:**
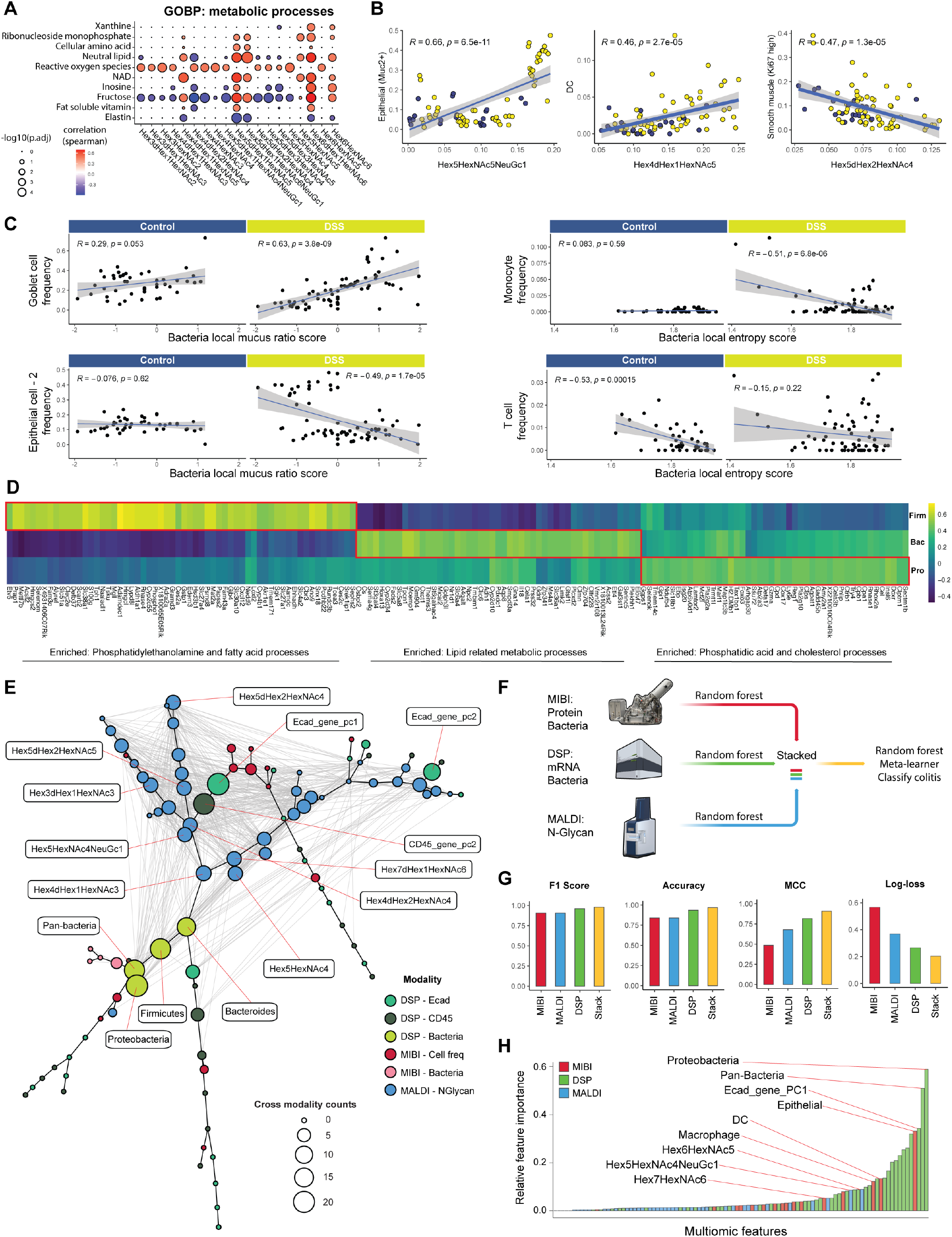
**(A)** Correlation between metabolic pathways scores from MicroCart-DSP and N-glycan levels from MALDI-MSI, based on aligned FOVs from adjacent slides. Metabolic pathways (from GO:BP database) or N-glycans with at least one significant (p.adj < 0.05) correlation were shown in the plot. **(B)** Dot plot showing correlations between cell type frequencies from MIBI and N-glycan levels from MALDI, based on the aligned FOVs from adjacent slides. Only relationships with significant (p.adj < 0.05, F test) correlations were shown. Color of dots indicates colitis status of the FOV. Line and shade indicate the linear relationship with 0.95 CI. **(C)** Dot plot showing correlations between cell type frequencies from MIBI and microbiome spatial metrics in the fecal regions from the same FOV. Left: cell frequencies and bacteria local mucus ratio score. Right: cell frequencies and bacteria local entropy score. Line and shade indicate the linear relationship with 0.95 CI. **(D)** Correlation (Z-normalized) between bacteria signals (from MicroCart-DSP) and host transcriptome signals from paired E-Cad compartments (adjacent to bacteria regions on the same tissue). The top 50 correlated genes per bacteria phylum (Firmicutes, Bacteroidetes, Proteobacteria) were shown in the heatmap. Annotation of the gene pathway was performed using Gene Ontology analysis with the top 50 correlated genes for each bacteria phylum. **(E)** Correlation network of the features across three different modalities. Each node represents a different feature, with color representing the modality, and the size of the node representing the number of significant (p.adj < 0.05) correlating cross-modality features it has. The backbone edges (black) and the layout of the nodes were generated by implementing a minimum spanning tree using the correlation-based distance among features. Gray edges between nodes indicate a significant (p.adj < 0.05) correlation between them. **(F)** Illustration of the schematics for training a tri-modality stacked ensemble prediction model. **(G)** Performance of the tri-modality stacked ensemble prediction model. F1 score, Accuracy, MCC (Matthews Correlation Coefficient) higher indicates better performance; Log loss (cross-entropy loss) lower indicates better performance. **(H)** Relative feature importance scores from the random forest classifiers. Higher value indicates higher contribution of the feature to the model prediction ability. Color indicates the modality type. Top 3 most important features from each modality type were labeled.

### Multi-omics spatial analysis of macrophage in colitis

Our observations thus far suggested the increased infiltration of monocytes and macrophages in the large intestine during colitis from both MicroCart-MIBI and -DSP adjacent sections (**Figs. 3D &4D**). The spatial multi-omics data generated here prompted us to focus our subsequent integrative analysis on tissue regions with high macrophage and monocyte infiltration in the large intestine. To realize a more comprehensive understanding of the cellular pathways and functional impacts of these tissue processes. We observed that macrophage and monocytes occupy spatially stratified niches around the smooth muscle cells in the intestinal muscular layer (**Fig. 6A**), suggestive of an orchestrated and spatially localized immunological response during colitis. We next performed a distance-based analysis to quantify our findings of infiltrating macrophages and monocytes into the muscle layer of the large intestine (**Supp. Fig. 8A**). Further analysis of the CD45 regions from MicroCart-DSP data also yielded significantly more collagen-related genes (*Col1a1, Col1a2, Col3a1, Col18a1, Col4a1*, and *Col5a1*) in addition to monocyte-linked genes (*S100a8, S100a11*, and *Ly6c1*), supporting our model for the increased proximity and infiltration of macrophages and monocytes into the intestinal smooth muscle cells (**Supp. Fig. 8B**). We further observed within the high macrophage and monocyte infiltration FOVs, elevated smooth muscle proliferation and migration-related gene signatures (**Fig. 6B, top**) and increased macrophage and monocyte chemotaxis-related gene signature (**Fig. 6B, bottom**) when compared to all other large intestine regions. These results were additionally supported by the further diminutive signatures observed for smooth muscle proliferation, migration, and macrophage and monocyte chemotaxis when compared across more stratified conditions (**Supp. Fig. 9A**). We reasoned that the infiltration of the macrophage and monocyte populations into these regions can initiate further downstream immune interactions and remodeling in colitis, and confirmed our hypothesis by modeling the pairwise cell type interactions using a permutation test (39, 52). Our results are indicative of an increased interaction between macrophages and monocytes with most cell types identified with MicroCart-MIBI (**Fig. 6C, Supp. Fig. 9B**).

To identify immune cell pathways and states associated with this functionality shift in macrophage and monocyte-infiltrated regions, we first utilized GSEA to confirm the upregulation of macrophage activities within the CD45+ immune compartments. Within the paired E-Cadherin+ epithelial regions, we observed the activation of multiple pathways specifically related to bacteria composition and microbial-linked immune suppression, including metal ion sequestering, humoral immune response to microbes, and hydrogen peroxide secretion. These results support a model in which macrophage and monocyte infiltration acts as a front-line host defense mechanism against bacterial components that breached the physical epithelium barrier during colitis (**Fig. 6D**).

We next investigated the relationship between gene programs within paired CD45+ and Ecad+ tissue regions for a systems-level understanding of immune-epithelial interactions in response to microbial infiltration. We first identified diverse gene programs using a consensus Non-negative Matrix factorization (cNMF) approach (53) from our MicroCart-DSP data, and performed a correlation analysis of these gene programs between the paired immune and epithelial regions. Interestingly, compared to other large intestinal regions, regions with high amounts of macrophage and monocyte infiltration (as detected via MIBI) exhibited more aggregated correlation gene program ‘hotspots’ (**Fig. 6E**), indicative of orchestrated immunological responses on the spatial level. We further investigated the functionality of these identified correlation hotspots of Ecad and CD45 regions by performing gene ontology analysis on the top contributing genes for these gene programs enriched within the hotspots. Our results identified pathways related to immune responses, immune cell population chemotaxis, phagocytosis and cell clearance (**Fig. 6F**), indicating that the macrophage and monocyte infiltration is a node for the diverse immune-epithelial tissue crosstalk during colitis in mouse intestine.

### Spatial tri-modal integration for a systems level analysis of colitis

The tri-modal MicroCart spatial-omics data generated, specifically 1) MIBI: cell phenotype, microbial composition and frequency information, 2) DSP: compartment-specific whole transcriptome and microbial quantification, and 3) MALDI-MSI: N-glycan levels. Given the varying scale of the data (**Fig. 1B**), we first manually aligned the tri-modal data on the individual FOV level, to maximize the concordance for downstream multi-omic investigation of the colitis samples. Given the link between glycan branching pathways and metabolite flux (54), our initial investigation focused on the correlations between pathway enrichment scores of various metabolic pathways and distinct glycan expression levels within the tissue regions. We first identified key metabolic pathways that are important for N-glycan expression, including those related to fructose, inosine, and NAD (**Fig. 7A**). We additionally link oxidative stress to changes of glycan expressions (55) (**Fig. 7A**). Further investigations identified statistically significant positive correlations between epithelium cell and dendritic cell frequencies and the glycans Hex5HexNAc5NeuGc1 and Hex4dHex1HexNAc5, and significant negative correlations between proliferating smooth muscle cell frequencies and Hex5dHex2HexNAc4 (**Fig. 7B**), suggesting the recruitment and depletion of key cell type-specific factors and their glycosylation states in the intestinal tissues.

We next investigated host responses to microbial infiltration during DSS-induced colitis. We observed specifically in colitis tissues, a positive correlation between goblet cell frequency and the bacteria local mucus ratio (**Fig. 7C, top left**), and a negative correlation with MUC2+ epithelial cells (**Fig. 7C, bottom left**), with the local mucus ratio as defined previously as the intermixing of bacteria and mucosal signals in the MIBI (**Fig. 3F**). We observed no significant corresponding relationship in the healthy control samples (**Fig. 7C, left**). We also observed negative correlations between monocyte frequencies with local bacteria Shannon entropy specific to colitis, and T cell frequencies, with the local bacteria entropy specific for healthy controls (**Fig. 7C, right**). Our results highlight a structured host response to local microbial perturbations, which prompted our next investigation into the linkage between bacteria and host spatial transcriptomic changes. We observed clear metabolic signatures as a result of the distinctive bacterial phylum composition (**Fig. 7D**), including for Firmicutes with phosphatidylethanolamine and fatty acid processes, Bacteroidetes with lipid-related metabolic processes, and Proteobacteria with phosphatidic acid and cholesterol processes compartmentalized to in the non-immune E-Cad+ regions (**Fig. 7D**). These results further solidify the host-microbial interactions and downstream effects post DSS-perturbation.

To gain systems-level perspective on colitis across all three spatial-omics modalities, we performed correlation network analysis encompassing all the features measured (56) (**Fig. 7E**). We identified key hubs in the microbial-host compartmentalized interactions within intestinal biology. Our constructed correlation network graph incorporated features representing cell population frequencies and bacteria signal strength (MicroCart-MIBI), singular value decomposition (SVD) dimension reduced transcriptomic principle components (MicroCart-DSP, to reduce feature numbers, **Supp. Table 7**), and N-glycan expression (MALDI-MSI). In the graph, nodes represent features, and the backbone edges (black) represent distances between features based on Spearman’s correlation and constructed via Minimal Spanning Tree (MST). Additional edges (gray) linking two nodes indicate significant correlations between the pair of features. Through this network, we identified key global features and significant correlations across modalities, indicative of the multitude of cell types, cell states, signaling pathways, and glycosylation patterns linked to an orchestrated host immunological response to microbial infiltration in the intestinal system. Our highlighted key signatures include bacteria signals, varying transcriptomic signatures from E-Cad and CD45 compartments, and varying N-glycans including Hex5dHex2HexNGc4, Hex5dHex2HexNGc5, Hex3dHex1HexNGc3.

We next sought to perform prediction of colitis status, using a stacked ensemble machine learning model, In line with previous demonstrations on the effectiveness of ensemble learning when applied to multi-omics data (57, 58). We first trained three individual random forest classifiers for each modality (spatial protein, RNA and glycans) to predict colitis status. We next applied a random forest ensemble learning layer on these three individual classifiers for the final prediction (**Fig. 7F**). Our results support the multi-omics ensemble learner as the highest performing model for classifying colitis status, when compared to single-modality classifiers (**Fig. 7G**). These results support the notion that a multitude of biomolecules (including proteins, RNA and glycans) are involved in the orchestrated immunological responses to diseases, as exemplified by colitis here. To identify key features for future hypothesis generation in colitis research, we tabulated the importance scores of cross-modality features within our classifier model (**Fig. 7H**). Notable high-importance features in our colitis classifier include microbial signatures (e.g., Proteobacteria), gene expressions in E-Cad compartments, and cell frequencies of Epithelial, DCs, and macrophages (**Fig. 7H**). These components warrant future detailed investigations to better understand the orchestrated tissue responses to microbial invasion of the gastrointestinal tract.

## Discussion

Here we introduce MicroCart, an approach for the integrative analysis of host-microbiome interactions through a spatialmulti-omics lens. Within our MicroCart framework, we provide a computational pipeline for the rapid design and validation of highly specific oligonucleotide probes targeting various components of the human microbiome, and present an efficient protocol enabling the preservation and simultaneous detection of bacterial, mucosal, and host signals *in situ*. This protocol is adaptable to diverse spatial-omics platforms, not limited to the MIBI and GeoMx-DSP as exemplified here. Enabled by the MicroCart pipeline, we designed and validated 16S rRNA targeting probes for distinct bacterial groups within the human gut microbiome, and assessed their on-target specificity. We next delved into exploring both systemic and localized shifts in the microbiome and host responses within mouse colitis models induced by DSS administration. The modularity of the MicroCart pipeline allowed our analysis to include MIBI (spatial proteomics), GeoMx-DSP (spatial transcriptomics), and MALDI (spatial glycomics) on adjacent sections of mouse intestinal tissues. Our findings revealed significant cellular compositional changes, transcriptomic responses related to various orchestrated immunological responses, and glycomics structural alterations during colitis. We further identified tissue-level remodeling interactions between host immune and epithelial cells in response to microbial infiltration, and pinpointed the pivotal role of macrophages as a key orchestrator in this dynamic process. We finally established a comprehensive trimodality feature network, employing machine learning approaches to identify key contributors to the status of mouse colitis. Our results highlight the need to understand the precise native tissue context of diseases and their constituents for future mechanistic and therapeutic work. In summary, MicroCart provides a powerful tool that also lays the foundation for future investigations seeking to unravel intricate cell-cell and cell-microbial interactions within the complex milieu of bacteria-host tissue environments.

## Materials & Methods

### Human 16S rRNA sequence pool construction

A comprehensive sequence pool for 16S rRNA sequences of intestinal microbiota was constructed following the method described in (28) with some modifications. Initially, sequences were obtained from the National Center for Biotechnology Information (NCBI) using the command `((“Homo sapiens”[Organism] OR Human[All Fields]) AND (intestinal[All Fields] OR gut[All Fields]) AND 16S[All Fields]) AND` `(“bacteria”[porgn] OR “archaea”[porgn])` `AND 1000:2000[SLEN]` in May 2019. A total of 79,223 sequences were collected. These sequences were then matched against the SILVA ribosomal database `SILVA_132_SSURef_Nr99_tax_silva_DNA.fasta`using Usearch (32) with the usearch_global command `-id 0.99 -strand plus -maxaccepts 1`. The matched sequences were then extracted from the SILVA database, and sequences shorter than 1.3 kb were filtered 674out. The final resulting sequence pool consisted of 12,936 near-full-length intestinal 16S rRNA sequences.

### Taxonomy assignment

The curated and length-filtered sequences from the sequence pool were assigned taxonomy information using the `Dada2` package in R (59). The `assignTaxonomy` function was used with the reference 680 database `hitdb_dada2`. A total of 4,881 sequences were assigned to the species level. To further enhance the taxonomy assignment, sequences that were not assigned to the genus level by `Dada2` but had genus information in SILVA were selected: If the family information for the sequence was 685 the same as `Dada2` and SILVA, the SILVA genus was then assigned to the sequence. The same criteria were applied to the species level. Consequently, 5,187 sequences were assigned to the species level. Sequences that still did not have species-level information were annotated with a “likely” taxonomy assignment. These sequences were searched against the sequences assigned with species information in the pool, using Usearch at >97% identity. Sequences with a best match >97% remained “Unknown” at the species level but were assigned a “likely” species based on the matched sequence. This type of annotation was not used during probe design, but the mismatch count of candidate probes on those sequences would be ignored if the probe targeted the annotated “likely” taxonomy group. The same process was performed at the genus level but with matching at >95% identity.

### Bacteria probe design

Fasta files containing the sequences and taxonomy information were loaded into ARB for probe design (29). The target group was selected, and the desired covering percentage and out-group hitting counts were provided. The length of the candidate probes was set to 15-30 bp. The candidate lists (`.prb` file) generated from ARB were then imported into R. Subsequently, the probes were screened following a similar approach as described in Wright et al. (30). Specifically, the melting temperature, secondary structure, and predicted hybridization efficiency were calculated for each candidate probe based on the experimental conditions (hybridization temperature, formamide concentration, and salt concentration) using a modified version of the function in R package `Decipher` (detail can be found in github repository). In this study, the input parameters were set as follows: hybridization temperature of 46 °C, formamide concentration of 40%, and sodium concentration of 390 mM. Candidate probes with a melting temperature (Tm) >60 °C, hybridization efficiency >0.8, and deltaG >-1.5 were selected as potential candidate probes.

For combinatorial probe design, the candidate probes produced from the previous step were utilized. Three individual probes from different regions of the 16S rRNA, as defined by ARB, were randomly chosen. These probe sets were matched against the sequence pool using the Usearch command `usearch_global` with an identity of 100%. The overall coverage (good hit) of the target group and the overall out-group hitting (bad hit) were recorded for each probe set. From a total of 2000 random combination trials, the optimized probe set with the desired coverage and out-group sensitivity was selected. To validate the specificity and coverage of the probes among abundant species in the human microbiome, the top 20 abundant human microbiome species in the phyla Firmicutes, Bacteroidetes, and Proteobacteria were acquired from the Human Microbiome Project (1). The ranking of the overall counts of the Operational Taxonomic Unit (OTU) table among 147 human stool samples, obtained through 16S rRNA sequencing, was used to determine the abundance. OTUs belonging to unclassified species were excluded. Subsequently, the full-length 16S rRNA sequences of the abundant species were retrieved from the SILVA database. The Usearch tool (`usearch_global: -id` 2 `1, -strand plus`) was employed to align the probes against the 16S rRNA sequences of the abundant species individually (**Supp. Table 8 & 9**). For future convenience, an R package has been compiled to facilitate the easy utilization of the functions described above, under github.

For probes designed for the GeoMx DSP platform, the same process described above was implemented, except the minimal probe length was set to 30 bps. For candidate probes that passed all requirements, a poly-A tail was added to the end of the probe to make the probe length of 35 bps, if not already larger or equal to that length.

After probe designing, probes (barcoded, fluorescent, or hapten versions) were either purchased from the Stanford PAN facility or from Integrated DNA Technologies (USA). Probes for the Nanostring GeoMx DSP platform were custom-designed (see above) and synthesized in collaboration with Nanostring (USA). The detailed sequences of these probes used in this manuscript can be found in the supplementary information (**Supplementary Table 2, 3, 5**).

### Bacteria strain culture

Bacteria strains used are either purchased from ATCC (Lactobacillus acidophilus ATCC® 4356™, Lactobacillus paracasei ATCC® BAA-52™, Lactobacillus reuteri ATCC® 23272™, Lactococcus lactis ATCC® 19435™, Bifidobacterium breve ATCC® 15700™, Bifidobacterium longum subsp. longum ATCC® 15707™, Clostridium perfringens ATCC® 13124™, Ruminococcus gnavus 35913™) or obtained from the Sonnenburg lab (Bacteroides fragilis NCTC 9343, Bacteroides finegoldii DSM 17565, Bacteroides dorei DSM17855, Bacteroides ovatus ATCC 8483, Lactobacillus plantarum ATCC BAA-793, Lactobacillus rhamnosus ATCC 53103).

Lactobacilli MRS Agar plate (Hardy Diagnostic, USA) was used to culture L. acidophilus, L. paracasei, L. reuteri, L. plantarum, L. rhamnosus, L. lactis. Blood Agar plate (Hardy Diagnostic, USA) was used to culture B. breve, B. longum,C. perfringens, R. gnavus. BHI agar plate (Hardy Diagnostic, USA) was used to culture B. finegoldii, B. dorei, B. ovatus, B. fragilis.

Each bacterial strain was inoculated onto the respective culture plates under sterile conditions. The plates were then placed in an air-tight gas pouch containing one pack of BD Difco™ GasPak™ EZ Gas Generating System (ThermoFisher B260683, USA) to create an anaerobic environment. The pouches were incubated at 37 °C for 48 hrs until harvest, allowing the bacteria to grow and develop.

### Bacteria MicroArray (MFPE) preparation for probe validation

To harvest the bacteria culture mentioned above, each plate was scraped using a sterile 20 µl pipette tip into a 1.5 ml Eppendorf tube filled with 1 ml of sterile 1x PBS. The tube was centrifuged at 800 xg for 10 mins, and the supernatant was discarded. The bacterial pellet was then incubated with 1 ml of Methacarn fixation solution (60% methanol, 30% chloroform, and 10% glacial acetic acid) for 30 mins at room temperature. During incubation, the tube was placed on a Mix Rack (ELMI, USA) and rotated at 10 rpm. After incubation, the bacteria were centrifuged at 800 xg for 10 mins, and the supernatant was discarded. The bacterial pellet was washed twice with 1 ml of PBS, each time centrifuging at 800 xg for 10 mins, and removing the supernatant. Subsequently, the bacteria were fixed with freshly prepared 4% PFA in 1x PBS for either 30 mins (for fluorescence / DSP imaging) or 6 hrs (for MIBI imaging). After fixation, the bacteria were washed twice with 1x PBS, each time centrifuging at 800 xg for 10 mins, and removing the supernatant. To facilitate embedding and storage, 20-50 µl of melted HistoGel (ThermoFisher, USA) was added to each bacterial sample. The sample was cooled at room temperature for 15 mins, followed by the addition of another 20-50 µl of melted HistoGel on top for sealing. The samples were then cooled at 4 °C for 1 hr until the HistoGel solidified. The solidified samples were removed from the Eppendorf tubes and placed in 9-compartment biopsy cassettes (EMS, USA). They were stored in 70% ethanol until processed in the pathology core at Stanford University, where they were embedded in paraffin blocks and sectioned into slides. Glass slides were used for fluorescence / DSP imaging, while gold slides (Ionpath, USA) were used for MIBI imaging. The slides were stored in vacuum chambers until they were ready for analysis. The fixation method used in this study, combining Methacarn and formalin fixation followed by paraffin embedding, was referred to as MFPE (Methacarn and Formalin-fixed, Paraffin-Embedded). This method enabled the preservation of mucus structure and protein epitopes.

### Mouse colitis model and tissue (MFPE) collection

C57BL/6J female mice that were 6 weeks old were obtained from Jackson Laboratory. To induce colitis, mice were provided with drinking water containing 3.5% dextran sulfate sodium salt (colitis grade, MPbio, USA) for a duration of 6 days. At the conclusion of the experimental period, mice were euthanized by CO_2_ asphyxiation. Two types of mouse intestinal tissues were collected: the distal 4 cm of the small intestine and large intestine parts containing formed fecal pellets. The collected tissues were placed in tissue cassettes and immediately immersed in methacarn solution (60% methanol, 30% chloroform, and 10% glacial acetic acid) for fixation, at RT for 3 hrs. Following methacarn fixation, the tissues were washed twice with 1x PBS for 10 mins each. Subsequently, the tissues were transferred to 4% PFA and fixed for 20 hrs. After fixation, the tissues were stored in 70% ethanol until further processing for paraffin embedding and sectioning into slides by the Stanford Pathology Core. Glass slides were used for sections intended for fluorescence/DSP/MALDI imaging, while gold slides (Ionpath, USA) were utilized for imaging by MIBI. All slides were stored in vacuum chambers until ready for use. All mice were maintained according to practices prescribed by the NIH at Stanford’s Research Animal Facility and by the Institutional Animal Care and Use Committee (IACUC, protocol 33699) at Stanford.

### Antibody conjugation for MIBI

The antibody conjugation process was conducted following a previously described protocol (60) utilizing the Maxpar X8 Multi Metal Labeling Kit (Fluidigm, USA). Initially, 100 µg of BSA-free antibody was subjected to washing using the conjugation buffer. Subsequently, the antibody was reduced by incubating it with a final concentration of 4 µM TCEP (Thermo Fisher Scientific, USA) for 30 mins in a water bath maintained at 37 °C. Following reduction, the antibody was mixed with Lanthanide-loaded polymers and incubated for 1.5 hrs in a water bath at 37 °C. Subsequently, the conjugated antibody was subjected to four washes using an Amicon Ultra filter (Millipore Sigma, USA). The resulting conjugated antibody was quantified using a NanoDrop spectrophotometer (Thermo Scientific, USA) in IgG mode, specifically measuring absorbance at 280 nm (A280). To ensure stability and preserve the conjugated antibody, the final concentration was adjusted using at least 30% (v/v) Candor Antibody Stabilizer (Thermo Fisher Scientific, USA). The conjugated antibody was then stored at 4 °C until further use. Information about antibody panels can be found in **Supplementary Table 4**.

### FISH hybridization on MFPE fixed samples

Fluorescence In Situ Hybridization (FISH) was conducted to validate the designed bacteria probes using fluorescent microscopy. Two types of samples were subjected to FISH: bacteria pellets (obtained from Methacarn + 4% PFA fixation for 30 mins) and mouse tissue sections (obtained from Methacarn + 4% PFA fixation for 20 hrs). The following steps were performed for FISH. For bacteria pellets, slides were initially baked at 70 °C for 15 mins, followed by two washes in xylene for 5 mins each. Subsequently, the slides were washed twice with 99.5% ethanol for 5 mins each. A Hydrophobic Barrier PAP Pen (Vector Labs, USA) was utilized to circle out the hybridization area. Rehydration of the samples was achieved by washing them with 2x SSCT for 5 mins. Hybridization was then performed using either primary probes directly labeled with fluorophore, or primary probes with secondary oligo barcodes. The hybridization buffer consisted of 2x SSCT, 10% Dextran sulfate, 1x Denhardt’s Solution, 40% Formamide, 0.01% SDS, 200 µg/ml Salmon sperm DNA, and oligonucleotide probes at concentrations ranging from 1 to 5 µM (1 µM for primary probes with fluorophore, and 2-5 µM for primary probes detected by secondary barcodes). The slides were incubated at 46 °C for 3 hrs in a humidity chamber. Following incubation, the hybridization buffer was removed, and the slides were subjected to three washes with 40% formamide in 2x SSCT at 46 °C, each lasting 10 mins. For samples stained with probes directly conjugated to fluorophores, the slides were quickly washed with 2x SSCT, stained with Hoechst 33342, mounted using Pro-Long™ Diamond Antifade Mountant, and sealed for imaging. In the case of samples stained with probes with secondary oligo barcodes, the slides were quickly washed with 2x SSCT and subjected to secondary detection probe staining in secondary hybridization buffer, containing 2x SSCT, 30% formamide, and 0.3 µM secondary detection probes labeled with fluorophores. The secondary hybridization was performed at room temperature for 20 mins. Subsequently, the slides were washed twice with 30% formamide in 2x SSCT for 5 mins each. Finally, the slides were quickly washed with 2x SSCT, stained with Hoechst 33342, mounted using Pro-Long™ Diamond Antifade Mountant, and sealed for imaging. Fluorescent images were acquired using a BZ-X710 inverted fluorescence microscope (Keyence) equipped with a CFI Plan Apo l 20x/0.75 objective (Nikon). To ensure accuracy during probe specificity validation experiment on bacteria pellets, the exposure times for each channel was set consistent: Hoechst (Hi-resolution setting) 1/25s; Cy3 (Hiresolution setting) 1/3s; Cy5 (Hi-resolution setting) 1/1.5s on the Keyence microscope.

### MicroCart staining for MIBI imaging

Gold slides with sections from bacteria pellets (Methacarn + 4%PFA 6 hrs) or tissue (Methacarn + 4%PFA 20 hrs) were baked at 70 °C for 30 mins, and then washed in xylene for 2 times, each 5 mins. Standard deparaffinization was performed thereafter (3x Xylene, 2x 100% EtOH, 2x 95% EtOH, 1x 80% EtOH, 1x 70% EtOH, 3x ddH_2_O; 1 min each). Epitope retrieval was then performed at 95 °C for 10 min at pH 9 with Dako Target Retrieval Solution (Agilent, USA), in a Lab Vision PT Module (Thermo Fisher Scientific). Slides were cooled to 65 °C and then removed from the PT Module, then cooled further in RT for 20 mins. After antigen retrieval, a Hydrophobic Barrier PAP Pen (Vector Labs, USA) was used to draw out the hybridization area. Slides were incubated in 2x SSCT (300 mM Sodium chloride, 30 mM Trisodium citrate, 0.1% (v/v) Tween-20) for 10 mins, then added with the hybridization buffer (2x SSCT, 10% Dextran sulfate, 1x Denhardt’s Solution, 40% Formamide, 0.01% SDS, 200 µg/ml Salmon sperm DNA, hapten oligo probes 1 - 3 µM each). Slides were then incubated at 46 °C for 3 hrs in a humidity chamber. After incubation, the hybridization buffer was removed, and slides were subject to three times washing with 40% formamide 2x SSCT at 46 °C, 10 mins each. Subsequently, slides went through a standard MIBI antibody staining process described before (61). Briefly: slides were quickly rinsed in MIBI Wash Buffer (1x TBS-T, 0.1% BSA) for 2 mins, and then blocked by Antibody Blocking Buffer (5% Donkey Serum, 0.05% NaN3 in 1x TBS-T) for 1hr at room temperature. Then slides were stained at 4 °C in an antibody cocktail (metal-conjugated) overnight. Slides were then washed twice by MIBI Wash Buffer, each 5 mins, then post-fixed by 4% PFA and 2% GA in 1x TBS-T for 15 mins. At last, slides were washed three times with 100 mM ammonium acetate, each 5 mins, air-dried, and stored in a vacuum chamber until MIBI imaging.

### MIBI imaging, processing, and analysis

Multiplexed imaging was conducted using a commercial MIBI-TOF mass spectrometer (MIBIscope™ System), equipped with a Xenon ion source. Running parameters on the instrument followed standard MIBI-TOF protocols (FOV size: 400 µm, Resolution setting: Fine 1ms, Depth: 1 layer). Subsequent to data acquisition, image processing was performed using custom code deposited on GitHub. In brief, metal counts were extracted from raw MIBI data files, and compensated for spectrum contamination using the methods described by the Toffy method (cite github). Then for mouse colitis samples, each individual image was manually separated into two masked regions: one containing fecal regions and the other containing luminal regions. The masks were manually drawn using FIJI (ImageJ). The images masked by the luminal mask underwent whole cell segmentation using Mesmer (62), where the dsDNA signal served as the nuclear channel and a linear summation of CD45, tubulin, and E-cadherin channels served as the membrane channel. Signal normalization was then performed within each MIBI run, whereby the median dsDNA intensities per segmented single cell from each field of view (FOV) were calculated. Subsequently, the signal intensity of all channels from each FOV were scaled up based on the ratio of the largest dsDNA median to the current FOV’s dsDNA median (within each MIBI run). The subsequent analysis diverged into two directions: 1) Analysis for host cells (within luminal masks): Signal aggregates were removed from images using empty mass channels (`mass_163`) as masks. Counts from single cells in segmented MIBI images were then extracted based on the segmentation generated by MESMER.

Single cells with size of less than 50 pixels or more than 2000 pixels were filtered out. Counts were normalized by the function `log1p` in R. Immune cells (CD45 >= 1.15 pre-log1p normalized) and non-immune cells (CD45 < 1.15 pre-log1p normalized) were clustered separately: for immune cells, “B220”, “CD3e”, “CD4”, “CD11b”, “CD11c”, “CD68”, “F480”, “IgA”, “Ly6g” were used for clustering; for non-immune cells, “Ecad”, “Ki67”, “MUC2”, “PNAD.1”, “SMA”, “Tubulin”, “Vimentin”, “CD31” were used for clustering. Unsupervised clustering was performed by functions `FindNeighbors` and `FindClusters` from R package `Seurat`, and subsequently manually annotated for cell types. 2) Analysis for bacteria (within fecal masks): To calculate local bacteria spatial metrics, the bacteria-related channels were first binarized, then a sliding window method was implemented: a window of size 100 x 100 pixels (∼ 40 µm) with sliding steps of 10 pixels was used. Windows that have an overlap with the host mask for more than 50% of the window area were removed for downstream analysis. For mucus-bacteria ratio calculation, the ratio between the positive percentage of PAN-bacteria (all bacteria) signal and the positive percentage of MUC2 signal within each sliding window was calculated. The medium ratio values inside each MIBI FOV were used for downstream analysis. For entropy calculation, function `stats.entropy` in python package `scipy` was used, where input was the percentage of positive pixels of each bacterial channel inside each sliding window. The medium entropy values inside each MIBI FOV were used for downstream analysis.

For correlative analysis between host and bacteria: cell type frequencies within each FOV were calculated based on cell annotations described above; local bacteria spatial metrics were calculated as described above. Values from the same FOVs were used to calculate the Pearson correlations and test statistics, with function `stat_cor()` in R package `ggpubr`. Details of the process are deposited in the github repository.

### MicroCart staining for GeoMx-Digital Spatial Profiling (DSP)

The GeoMx-DSP mouse Whole Transcriptome Atlas (WTA) panel was stained as previously described but with modification to be compatible with MicroCart (13). In brief, adjacent glass slides from MIBI imaging slides were baked at 70 °C for 30 mins, and then washed in xylene for 2 times, each 5 mins. Standard deparaffinization was performed thereafter (3x Xylene, 2x 100% EtOH, 2x 95% EtOH, 1x 80% EtOH, 1x 70% EtOH, 3x ddH_2_O; 1 min each). Epitope retrieval was then performed at 95 °C for 10 min at pH 9 (Dako Target Retrieval Solution, S236784-2) in a Lab Vision PT Module (Thermo Fisher Scientific). Slides were cooled to 65 °C and then removed from the PT Module, then cooled further in RT for 20 mins. After antigen retrieval, a Hydrophobic Barrier PAP Pen (Vector Labs, USA) was used to draw out the hybridization area, and then washed 1 min in 1x PBS. Slides were then digested by Protease K (0.1 µg/ml) for 5 mins at 37 °C, and then washed with 1x PBS. For BMA samples (if used), the Protease K step was skipped. Subsequently, slides were fixed by 10% NBF for 5 min at room temperature, then the fixation process was stopped by 5 mins of 1x NBF Stop Buffer wash, followed by 5 mins 1x PBS wash. The slides were first stained with custom-made bacteria probes with DSP NGS barcodes (Nanostring), then stained with the mouse WTA panel. In detail: slides were first stained with custom bacteria NGS probes in hybridization buffer (2x SSCT, 10% Dextran sulfate, 1x Denhardt’s Solution, 40% Formamide, 0.01% SDS, 200 µg/ml Salmon sperm DNA,bacteria DSP probes 5 nM) at 46 °C for 3 hrs, in a humidity chamber. After incubation, the hybridization buffer was removed, and slides were subject to three times washing with 40% formamide 2x SSCT at 46 °C, 10 mins each, then followed with a quick 2x SSC wash. Afterwards, Nanostring DSP mouse WTA detection probes were then applied to the slides and incubated overnight (∼ 18 hrs) at 37 °C. After hybridization, slides were washed in Stringent Wash Buffer (2x SSC, 50% Formamide) 2 times, each 5 mins at RT. Slides were then washed by 2x SSC twice, 2 mins each. Buffer W was then applied to the slides for 30 mins, followed by antibody staining for 1hr 1:100 dilution of CD45-Alx647 (D3F8Q, CST), and 1:100 dilution of E-Cadherin-Alx594 (24E10, CST). Slides were then washed by 2x SSC twice, 5 mins each, and stained with 500 nM SYTO 13 for 15 mins, then loaded onto the GeoMx DSP machine. For the bacteria pellet sample, the process was the same as described above, but with two differences: 1), the antigen retrieval step was skipped. 2), 2x SSC, instead of the mouse WTA detection panel, was used during the first staining step, as no mouse cells were present in the bacteria pellet samples.

### Digital Spatial Profiling data acquisition and analysis

For the GeoMx DSP sample collection, we followed the guidelines provided in the GeoMx DSP instrument user manual (MAN-10088-03). The process involved selecting specific regions of interest (ROIs) that were imaged by MIBI on the adjacent gold slide. Three types of ROIs were chosen: 1) luminal regions positive for CD45, 2) luminal regions positive for Ecad, and 3) adjacent fecal regions. For bacteria pellet samples (validation), ROIs were circles with 100 µm radius with bacterial cells. Sample collection was performed according to the designated ROIs. Subsequently, the Nanostring NGS library preparation kit was utilized. Each collected ROI was uniquely indexed using Illumina’s i5 x i7 dual-indexing system. A PCR reaction was carried out with 4 µl of collected samples, 1 µM of i5 primer, 1 µM of i7 primer, and 1x Nanostring library prep PCR Master Mix. The PCR conditions included incubation at 37 °C for 30 mins, 50 °C for 10 mins, an initial denaturation at 95 °C for 3 min, followed by 18 cycles of denaturation at 95 °C for 15 s, annealing at 65 °C for 60 s, extension at 68 °C for 30 s, and a final extension at 68 °C for 5 mins. The PCR product was purified using two rounds of AMPure XP beads at a 1.2x bead-to-sample ratio. The libraries were then subjected to paired-end sequencing (2 x 75 bp) on a NextSeq550 platform (Novogene). The NGS barcodes from the Nanostring mouse WTA panel and custom bacteria probes were mapped and counted using the commercial GeoMx Data Analysis software pipeline, using FASTQ files generated from NGS sequencing. The resulting data underwent quality control and normalization steps, using the R package `Geomx-Tools` provided by Nanostring. Initially, ROIs and probes that did not meet the default quality control requirements were filtered out and excluded from subsequent analyses. Next, raw probe counts were transformed into a gene-level count matrix by calculating the geometric mean of the probes corresponding to each gene. Normalization of gene counts was performed using the ‘Q3 norm (75th percentile)’ method recommended by `Geomx-Tools`. The normalized gene counts (Q3 normed) were then used for downstream analyses. Differentially expressed genes (DEG) between control and DSS treated samples were identified using a linear mixed-effect model (LMM) documented by `Geomx-Tools`. Gene set enrichment analysis (GSEA) was performed with R package `GSEA` with function `gsea` and database ‘GO:BP’ (63). Details of the process are deposited in the github repository.

### MALDI-MSI N-Glycan data acquisition and analysis

The tissue preparation process was followed as previously described (64). In brief, glass slides with MFPE mouse intestinal tissues were baked at 70 °C for 30 mins, and then washed in xylene for 2 times, each 5 mins. Standard deparaffinization was performed thereafter (3x xylene, 2x 100% EtOH, 2x 95% EtOH, 1x 80% EtOH, 1x 70% EtOH, 3x ddH_2_O; 1 min each). Epitope retrieval was then performed at 95 °C for 10 min at pH 9 (Dako Target Retrieval Solution, S236784-2) in a Lab Vision PT Module (Thermo Fisher Scientific). Slides were cooled to 65 °C and then removed from the PT Module, then cooled further in RT for 20 mins. Afterwards, slides were dried overnight in a desiccator. Then, a total of 15 passes of the PNGase F PRIME enzyme at 0.1 µg/µl was applied to the tissue slides, at a rate 25 µl/min with a velocity of 1200 mm/min and a 3 mm offset at 10 psi and 45 °C using an M3+ Sprayer (HTX Technologies, USA). Enzyme-sprayed slides were then incubated in prewarmed humidity chambers for 2 hrs at 38.5 °C for deglycosylation. After incubation, a total of 14 passes of 7 mg/ml CHCA matrix in 50% ACN/0.1% TFA was applied to the deglycosylated slides at a rate of 70 µl/min with a velocity of 1300 mm/min and a 3 mm offset at 10 psi and 77 °C using the same sprayer. Washing steps using low and high-pH solutions and water were performed between enzyme and matrix applications to clear the sprayer headline. After matrix deposition, slides were desiccated until analysis. To assist batch effect correction for MALDI signals, 4 tissue cores from the same human TMA were sectioned into each glass slide with mouse intestine samples, and utilized as baseline normalizations for downstream analysis.

A timsTOF fleX MALDI-2 mass spectrometer (Bruker Daltonics, Germany) equipped with a 10 kHz SmartBeam threedimensional (3D) laser operating in positive mode with a spot size of 10 µm was used to detect released N-glycans at a high resolution. 200 laser shots per pixel over a mass range of 800 to 4000 m/z were collected for analysis, with an ion transfer time of 120 µs, a prepulse storage time of 28 µs, a collision frequency of 4000 Vpp, a multipole frequency of 1200 Vpp, and a collision cell energy of 10 eV.

Following MALDI data analysis, signals were extracted and generated into .tiff (per glass slide) images using https://github.com/angelolab/maldi-tools developed by the Angelo lab. To account for batch effects among different slides during N-Glycan level comparisons between colitis status, the signals from each N-Glycan molecule were normalized to the same scale, based on the average ratio calculated between the corresponding 4 control tissue cores (from Human TMA with muscle/epithelial tissues) across slides. To perform pixel-level clustering on N-Glycan signals, `pixie` was implemented as previously described (49).

Briefly, `harmony` (65) was performed first to further correct the batch effects at the latent space level. Pixel level N-glycan signals from each image were flattened into *n × p* dimensional matrix, where *p* is the number of N-Glycan, and *n* is the total number of pixels in each image/slide.

We then concatenated the vectors among all slides (*m × p*, from 4 slides in total), and utilized SVD to denoise and reduce the matrix to *m × k*, where *k* was set to 30. Subsequently the function `run_harmony` from python package `harmonypy` was used to run on the dot product of *U* and *D* matrix from the SVD process in the previous step, where the slide id was used as batch labels. The resulting harmony `Z_corr` loading matrix was then used to calculate the dot product with the V matrix from the SVD process in the previous step, and reformatted back to matrices with the same dimensions as the original .tiff image (eg. *h × w × p*). This resulted in a ‘harmony - corrected’ version of the image, and these images were used as input for the `pixie` pipeline. The number of pixel clusters to be defined was set to 20 clusters, and no Gaussian blurring was applied to images and other parameters were set as default for the pipeline. Details of the process are deposited in the github repository.

### Macrophage analysis

To identify high macrophage /monocyte infiltration tissue areas, MIBI FOVs were ranked by macrophage and monocyte (combined) percentage, and the top 15 FOVs (and the paired DSP regions) were labeled as high infiltration, and the rest of all FOVs were labeled as other. Gene pathway scores were calculated based on gene expression data from DSP-CD45 regions, using the function `gsva` in R package `GSVA`. For ‘Smooth muscle proliferation score’, genes from the GO:BP database term: ‘GOBP_SMOOTH_MUSCLE_CELL_PROLIFERATION’ were used. For ‘Smooth muscle migration score’, genes from the GO:BP database term: ‘GOBP_SMOOTH_MUSCLE_CELL_MIGRATION’ were used. For ‘Macrophage chemotaxis score’, genes from the GO:BP database term: ‘GOBP_MACROPHAGE_CHEMOTAXIS’ were used. For ‘Monocyte chemotaxis score’, genes from the GO:BP database term: ‘GOBP_MONOCYTE_CHEMOTAXIS’ were used. Cell-cell interaction analysis was performed as previously described (39, 52). In brief, for each individual macrophage or monocyte, the Delaunay triangulation for neighboring cells (within 50 µm) was calculated based on the XY position with the `deldir` R package. To establish a baseline distribution of the distances, cells were randomly assigned to existing XY positions, for 1000 permutations.

The baseline distribution of the distance was then compared to the observed distances using a Wilcoxon test (two-sided). The log2 fold enrichment of observed mean over expected mean for each interaction type was plotted for interactions with a p-value < 0.05. The test results also include the interactions in both directions (eg. Macrophage => T and T => Macrophage). GSEA analysis on the high infiltration regions was implemented similarly as described in previous sections.

Gene programs for DSP host WTA data were identified via cNMF as previously described (53). In brief, functions from python package `cnmf` were used on the top 8000 variable genes in the q3-normalized DSP gene expression data. The rank in cNMF (number of gene programs) was set to 35 (determined via function `k_selection_plot`). After identifying the gene programs, the Spearman correlation of the programs scores between paired CD45 and Ecad regions was calculated, and plotted as a heatmap with programs clustered by hierarchical clustering. To annotate each ‘correlation hotspot’ in the heatmap, the top 10 contributing genes for each gene program (identified from `gep_scores` from package `cnmf`) within the selected ‘hotspots’ were grouped, and Gene Ontology term enrichment analysis was performed on the grouped genes from each hotspot, using the function `enrichr` in R package `enrichR`, with database ‘GO_Biological_Process_2015’. Details of the process are deposited in the github repository.

### Multiomic analysis, correlation network, and stacked ensemble model

Multiomic information of MIBI, DSP, and MALDI from each individual intestinal tissue region were gathered for analysis. FOVs from MIBI and ROIs from DSP were paired and used for downstream analysis. For MALDI data, masks where the MIBI FOVs and DSP ROIs were acquired on the tissue were manually generated, and glycan expression profiles were extracted for each FOV. This process created a MIBI-DSP-MALDI tri-modality paired data across different tissue regions.

For correlative analysis between GO:BP-metabolic processes and glycan expression: gene terms with pattern ‘metabolic_processes’ were selected from the GO:BP database, and the corresponding genes for each gene term were extracted and used to calculate a gene term enrichment score for each tissue region by function `gsva` from R package `GSVA`. Gene terms of metabolic processes with the top 20 highest variation across samples (based on gsva scores) were selected, and the Spearman correlation between the glycan expressions and gene terms were calculated. For visualization purposes, features (gene terms and glycans) with at least one significant correlation (p.adjusted < 0.05) were shown in the heatmap. For correlative analysis between cell type frequencies and glycan expression: the correlation between cell frequencies within each tissue region from MIBI data and glycan expressions were calculated and plotted.

For correlative analysis between host transcriptome and bacteria signal, Spearman correlations were calculated between each mouse WTA gene inside ‘Ecad’ regions (large intestine), and bacteria signals (‘Firmicute’, ‘Bacteroidetes’, ‘Proteobacteria’) in the adjacent ‘Fecal’ regions. The top 50 genes with the highest absolute Spearman correlation values for each type of bacteria signal were used for plotting and analysis. GSEA was performed with R package `GSEA` with function `gsea` and database GO:BP on the highly correlative genes for each bacteria signal type. For correlative analysis between cell type frequencies and microbiome local spatial metrics, the Spearman correlations were calculated between within each tissue region from MIBI FOVs, and the adjacent fecal regions.

For cross-modality feature correlation network construction: For MIBI, host cell-type percentages and normalized bacteria signal from each FOV were used as input; For DSP, we implemented dimension reduction first to decrease the number of features. We applied SVD to denoise and reduce the gene expression matrix from CD45 and Ecad regions separately, with k set to 20, and the dot product of the U and D matrix were used as the reduced gene components. The loading values from the V matrix were used to identify the top 10 positive contributing (high loading value) and top 10 negative contributing (low loading value) genes for each gene component. The CD45 and Ecad gene components, along with bacteria signals from fecal regions from each ROI were used as input; For MALDI, N-Glycan signals from each manually aligned mask were used as input. Subsequently, features from these three modalities were concatenated, and a correlation (Spearman) matrix was calculated with function `rcorr.adjust` from R package `RcmdrMisc`. The correlation matrix was then transformed into a distance matrix to construct a graph with Minimal Spanning Tree (MST), with function `mst` from R package `ape` and function `graph.adjacency` from R package `igraph`. The MST graph was used as the backbone of the network. Subsequently, we constructed a second graph, where nodes (feature) were connected together by an edge, if there was significant (p.adj < 0.05) correlation observed. Finally, the graphs were plotted by functions from R packages `ggplot` and `ggnetwork`, where the node placements were determined by MST graph layout, and edges connected by either MST graph (backbone) or the second significant correlation graph.

For the stacked ensemble prediction model, single modality or stacked multi-omic features were used to classify colitis status. Single modality prediction models were Distributed Random Forest (DRF) classifiers with function `h2o.randomForest` from R package `h2o`; the stacked ensemble prediction model was achieved by stacking the 3 single modality DRF classifiers with function `h2o.stackedEnsemble`, using a DRF metalearner. All single or stacked models used the same 60% - 40% train test data split, 5-fold cross-validation with the same seed. Feature importance scores were calculated based on function `h2o.varimp`, where the importance percentages from each single modality model were extracted, and further weighted by drop-out ensemble model performances tests, in order to scale up features from more important modalities. Details of the process are deposited in the github repository.

## DATA AVAILABILITY

All data generated in this study will be deposited on Zenedo and freely accessible upon publication.

## CODE AVAILABILITY

Code related to 16S rRNA probe design: https://github.com/BokaiZhu/microbiomeFISH; Code related to analysis performed in this manuscript: https://github.com/BokaiZhu/microcart_analysis.

## ACKNOWLEDGEMENTS

The authors thank Justin Sonnenburg, Erica Sonnenburg, and members of the Sonnenburg lab for their extended help on this project. The authors thank members of Ionpath Inc., Nanostring Inc., and Bruker Inc. for their technical support. The authors thank the insightful discussion with lab members from G.P.N., S.J., A.S., M.A. labs. The authors thank the Stanford PAN facility and Pathology core (Pauline Chu). B.Z. was previously supported by the Stanford Graduate Fellowship during this work. G.K.G. is supported by NIGMS R35GM149270, BWH President’s Scholar Award, NIGMS R01GM130777, NSF MTM2 2025515. S.J. is supported by NIH DP2AI171139, P01AI177687, R01AI149672, a Gilead’s Research Scholars Program in Hematologic Malignancies, a Sanofi iAward, the Bill & Melinda Gates Foundation INV-002704, the Dye Family Foundation, and previously by the Leukemia Lymphoma Society Career Development Program. G.P.N. is supported by the Rachford and Carlota A. Harris Endowed Professorship. A.K.S. is supported by NIH 1P01AI177687-01, and Bill & Melinda Gates Foundation INV-055706.

This article reflects the views of the authors and should not be construed as representing the views or policies of the institutions that provided funding.

## AUTHOR CONTRIBUTIONS

Conceptualization: B.Z., S.J., G.P.N

Experiment: B.Z., Y.B., Y.Y., X.L., X.R.C., H.C.

Analysis: B.Z., Y.B., J.Y.

Contribution of reagents, tools, or technical expertise: G.k.G., E.S., J.S., M.A., A.k.S.

Supervision and funding: A.k.S., G.P.N., S.J.

